# The dynamic interplay between ATP/ADP levels and autophagy sustain neuronal migration *in vivo*

**DOI:** 10.1101/2020.04.02.022228

**Authors:** Bressan Cedric, Pecora Alessandra, Gagnon Dave, Snapyan Marina, Labrecque Simon, De Koninck Paul, Parent Martin, Saghatelyan Armen

## Abstract

Cell migration is a dynamic process that entails extensive protein synthesis and recycling, structural remodeling, and a considerable bioenergetic demand. Autophagy is one of the pathways that maintain cellular homeostasis. Time-lapse imaging of autophagosomes and ATP/ADP levels in migrating cells in the rostral migratory stream of mice revealed that decrease in ATP levels force cells into the stationary phase and induce autophagy. Genetic impairment of autophagy in neuroblasts using either inducible conditional mice or CRISPR/Cas9 gene editing decreased cell migration due to the longer duration of the stationary phase. Autophagy is modulated in response to migration-promoting and inhibiting molecular cues and is required for the recycling of focal adhesions. Our results show that autophagy and energy consumption act in concert in migrating cells to dynamically regulate the pace and periodicity of the migratory and stationary phases in order to sustain neuronal migration.

**Highlights:**

- ADP levels dynamically change during cell migration
- A decrease in ATP levels leads to cell pausing and autophagy induction via AMPK
- Autophagy is required to sustain neuronal migration by recycling focal adhesions
- Autophagy level is dynamically modulated by migration-promoting and inhibiting cues

## Introduction

Cell migration is a crucial mechanism for normal embryonic development, and a cell migration deficit leads to the loss of embryos or functional and cognitive impairments (Ayala et al., 2007). Although neuronal migration largely ceases in the postnatal period, it is preserved in several areas associated with postnatal neurogenesis such as the cerebellum, hippocampus, and subventricular zone (SVZ), which is the area associated with the production of neuronal precursors destined for the olfactory bulb (OB). Neuronal precursors born in the SVZ migrate tangentially along the rostral migratory stream (RMS) and, once in the OB, turn to migrate radially and individually out of the RMS into the bulbar layers (Kaneko et al., 2017; Gengatharan et al., 2016). In the postnatal RMS, neuroblasts travel in chains ensheathed by astrocytic processes along blood vessels (Kaneko et al., 2010; Snapyan et al., 2009; Whitman et al., 2009). Cell migration is a very dynamic process composed of migratory phases intercalated with stationary periods and is accompanied by structural remodeling, organelles dynamic and protein trafficking and turnover (Tanaka et al., 2017; Webb et al., 2002). Cell migration also entails a considerable bioenergetic demand, and it is still unclear how migrating cells regulate cellular homeostasis by clearing damaged organelles and aggregated/misfolded proteins, and how the metabolic requirements of neuroblasts are dynamically regulated during the different phases of cell migration.

Autophagy is a key evolutionarily conserved intracellular process that controls protein and organelle degradation and recycling. During this process, intracellular materials are sequestered inside double-membrane vesicles (autophagosomes) that then fuse with lysosomes to form autolysosomes in which aggregated/misfolded proteins and damaged organelles are degraded (Yang and Klionsky, 2010). Autophagy plays an important role in neurodevelopment (Mizushima and Levine, 2010) and alterations in autophagy have been observed in various human diseases (Yang and Klionsky, 2010). Autophagy is under control of a large protein complex and may be initiated by the Ulk1 and Ulk2 proteins, the downstream effectors of AMP kinase (AMPK) and mTOR (Petherick et al., 2015; Egan et al., 2011). The initiation of autophagy is accompanied by the formation of Atg5-Atg12 complexes and by Atg7-dependent conversion of microtubule-associated protein light chain 3 (LC3) from the inactive cytoplasmatic form into the membrane-bound lipidated form (LC3-II), which is found at the surface of autophagosomes (Simonsen and Tooze, 2009). While the role of autophagy in neuronal development is well understood (Mizushima and Levine, 2010), little is known about its role in neuronal migration. It has recently been shown that autophagy promotes cell migration (Kenific et al., 2016; Sharifi et al., 2016). However, other studies have shown that autophagy can have a negative impact on cell migration (Li et al., 2015; Tuloup-Minguez et al., 2013), raising a debate about the exact role played by autophagy in neuronal migration. Furthermore, most of our knowledge about the role of autophagy in cell migration come from studies on epithelial and tumor cell cultures (Kenific et al., 2016; Sharifi et al., 2016; Li et al., 2015; Tuloup-Minguez et al., 2013). As such, *in vivo* studies on the effect of autophagy on the migration of neuronal cells are required. It is also unclear how autophagy is dynamically regulated in migrating cells and how it is induced.

We show that autophagy is dynamically modulated in migrating cells in the RMS and that autophagy levels change in response to molecular cues that promote or inhibit migration. Changes in autophagy levels are linked to changes in the recycling of the focal adhesion protein paxillin. We used time-lapse imaging of autophagosomes and measurements of ATP/ADP levels in migrating cells to show that, under basal conditions, autophagy levels are correlated with the bioenergetic requirements of cells and that decreases in ATP/ADP levels during the migratory phases lead to the entry of cells into the stationary phase, the activation of AMPK, and the induction of autophagy. The genetic disruption of autophagy-related proteins and AMPK-downstream targets such as Ulk1, Ulk2, Atg5 and Atg12 hampers neuronal migration by prolonging the length of the stationary phase, leading to the accumulation of cells in the RMS. Our results show that autophagy, by sustaining neuronal migration, regulates the faithful arrival of neuronal precursors in the OB. Our results also mechanistically link autophagy and energy consumption in migrating cells to the dynamic regulation of the pace and periodicity of the migratory and stationary phases.

## Results

### Autophagy is dynamically regulated in migrating neuroblasts

To investigate the involvement of autophagy in cell migration *in vivo*, we first performed immunostaining for two key autophagic proteins, microtubule-associated protein 1 light chain 3 (LC3, isoforms A and B) and Atg5, on sagittal sections of the adult mice forebrain. Neuroblasts were labeled using GFP-expressing retroviruses injected into the SVZ. LC3 and Atg5 were highly expressed in GFP+ migrating neuroblasts in the RMS (**Figure 1A**). Between 55% and 60% of the neuroblasts were immunostained for Atg5 or LC3 (**Figure 1A-C**). In contrast, less than 10% of astrocytes in the RMS of GFAP-GFP mice were immunolabeled for autophagy-related proteins (**Figure 1A-C**). To confirm the presence of autophagosome vesicles, which are characterized by a double membrane (Klionsky et al., 2016), we examined RMS samples by electron microscopy. We took advantage of tamoxifen-inducible GlastCreERT2::GFP mice in which GFP is expressed in astrocytes and stem cells and their progeny following tamoxifen injection. GFP+ immunostained neuroblasts, which were identified based on their typical bipolar morphology with leading and trailing processes, exhibited numerous double-membrane autophagosomes (**Figure 1D)**. LC3 is present in a non-lipidated form (LC3-I) in the cytoplasm and a conjugated phosphatidylethanolamine lipidated form (LC3-II) in the autophagosomal membrane (Kabeya et al., 2004). The presence of LC3-II is a sign of autophagosome formation (Klionsky et al., 2016). Western blots of micro-dissected OB and RMS, two regions that contain migrating neuroblasts, showed the presence of autophagosome-associated LC3-II (**Figure 1E)**. The presence of autophagosomes in neuroblasts could be due to active autophagic flux or interrupted autophagy, such as the absence of fusion of autophagosomes with lysosomes, leading to vesicle accumulation. To distinguish between these two possibilities, we infected neuroblasts with a retrovirus encoding the RFP-GFP-LC3 fusion protein. The expression of RFP-GFP-LC3 makes it possible to distinguish autophagosomes (RFP+/GFP+ vesicles) from autolysosomes (RFP+ vesicles) due to GFP quenching in the acidic lysosomal environment. We observed a higher proportion of RFP+ than of GFP+/RFP+ vesicles and a highly dynamic bidirectional movement of RFP-LC3 puncta (**Figures 1F-I**; **Supplementary Movie 1**). Interestingly, 1-h time-lapse imaging of migrating neuroblasts infected with the RFP-GFP-LC3 retrovirus revealed a dynamic regulation and non-homogeneous distribution of autophagosomes/autolysosomes during the different cell migration phases (**Figure 1I**). The migratory phases of neuroblasts were associated with lower density of RFP+ puncta, whereas the entry of cells into stationary phases led to a higher density of autophagosomes/autolysosomes (**Figure 1I**). Altogether, these results suggest that autophagy is an active and uninterrupted process and that it is dynamically regulated during the different phases of cell migration.

**Figure 1:**
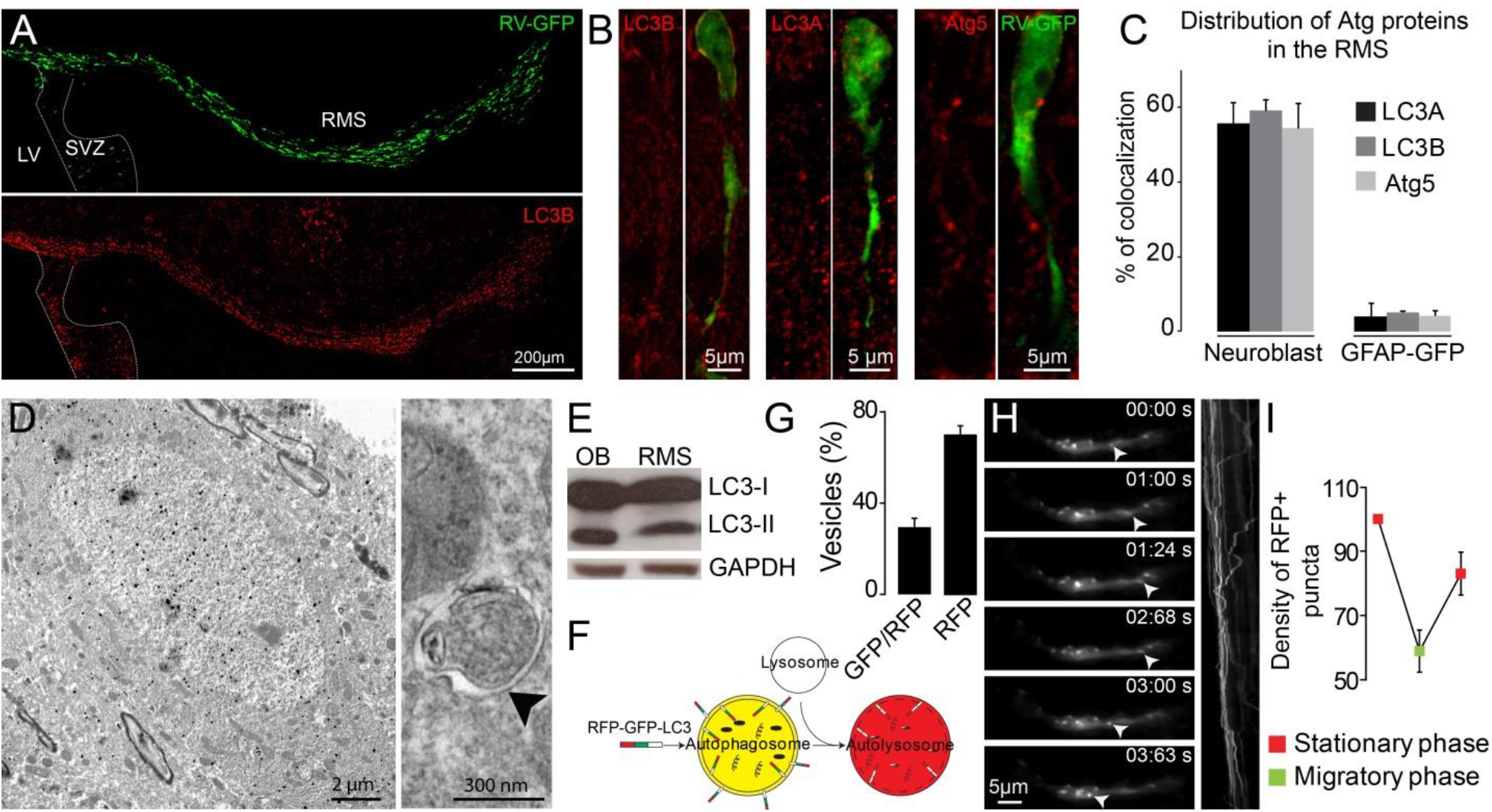
Migrating cells express autophagy-related proteins and display active autophagic flux. **A-C:** Confocal images and quantification showing Atg5, LC3A, and LC3B expression by neuroblasts and astrocytes. The percentage of colocalization was calculated as the number GFP+ cells (neuroblast or astrocytes) presenting labelling for one of these proteins over the total numbers of GFP+ cells. **D:** Electron microscopy (EM) analysis of autophagy in neuroblasts in the RMS of Glast-CreErt2xGFP mice. Neuroblasts are labeled with anti-GFP immunogold particles. An autophagosome is indicated with an arrowhead. **E:** Western blot analysis of LC3 in OB and RMS samples. **F:** Scheme of the RFP-GFP-LC3 fusion protein used to study autophagic flux. The RFP-GFP-LC3 fusion protein makes it possible to label autophagosomes in the RFP and GFP. Only autolysosomes were labelled in the RFP because GFP was quenched due to the acidic pH in the lysosomes. **G:** Percentage of GFP+/RFP+ vesicles versus RFP+ vesicles in neuroblasts of fixed brain sections, obtain after stereotaxic injections of retrovirus encoding RFP-GFP-LC3. **H:** Time-lapse imaging and kymograph of RFP-LC3 vesicles in migrating neuroblasts. **I:** Quantification of changes in the RFP+ puncta of individual neuroblasts during different phases of cell migration. Density of autophagosomes/autolysosomes in neuroblasts during the different migration phases is ploted. Data are expressed as means ± SEM. See also **Supplementary Movie 1**.

### The suppression of autophagy hampers cell migration and leads to the accumulation of neuroblasts in the RMS

To investigate the role of autophagy in cell migration, we first assessed migration of GFP-labeled neuroblasts in the RMS following application of bafilomycin (4 μM), which is known to inhibit the fusion of autophagosomes with lysosomes (Yamamoto et al., 1998). We stereotactically injected a GFP-encoding retrovirus in the SVZ and analyzed cell migration in the RMS 7-10 days later. Time-lapse imaging of migrating neuroblasts in acute brain sections revealed that bafilomycin reduced the distance of migration for GFP+ neuroblasts (46.1 ± 2.9 μm in control vs. 27.6 ± 3.3 μm following bafilomycin application; *p* < 0.001, n = 55 and 41 cells from 3 mice per condition, respectively; **Figure 2A-D**). This decrease was caused by a reduction in the percentage of the migratory phases (53.2 ± 1.6% and 33.8 ±1.8 μm/h in control and bafilomycin conditions, respectively; *p* < 0.001 **Figure 2F**), with no change in the speed of migration, which was estimated exclusively during the migratory phases (114.9 ± 5.0 μm/h and 97.6 ± 5.6 μm/h in control and bafilomycin conditions, respectively, **Figure 2E**). In these as well as all other migration experiments, we did not take into consideration cells that remains stationary during the entire imaging periods, which likely underestimate the autophagydependent effects on cell migration.

**Figure 2:**
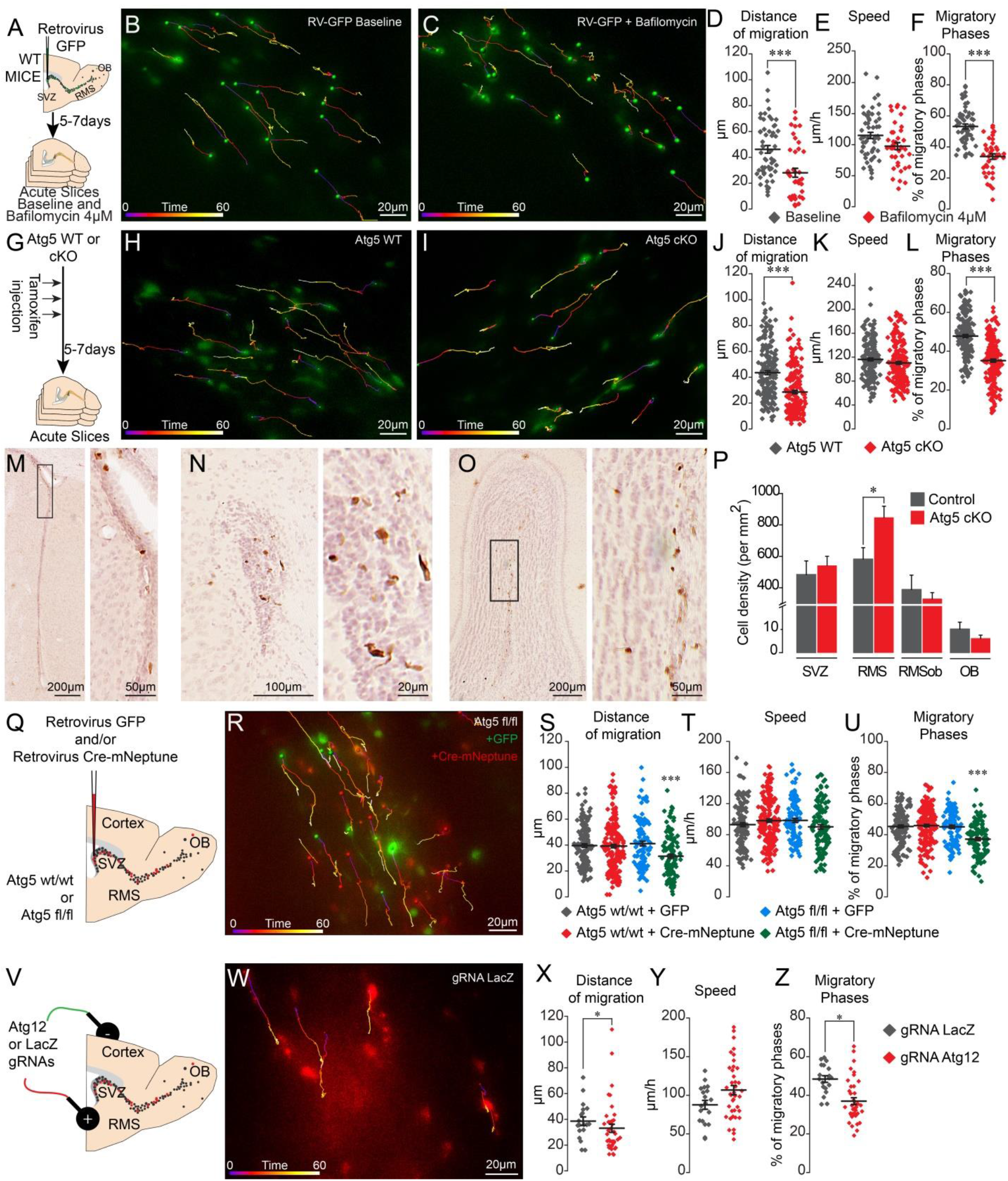
Pharmacological and genetic impairment of autophagy-related genes disrupts cell migration. **A:** Experimental design for pharmacological inhibition of autophagy using bafilomycin application. **B-C:** Time-lapse imaging of GFP-labeled neuroblasts in acute brain sections. **D-F:** Distance of migration, speed of migration, and percentage of migratory phases of neuroblasts under control condition and following application of bafilomycin (n = 55 and 41 cells from 3 mice for control and bafilomycin conditions, respectively). *** p < 0.001 with the Student t-test. **G:** Experimental design to conditionally delete Atg5 in adult mice. **H-I:** Time-lapse imaging of Atg5 cKO and WT neuroblasts in acute brain sections. **J-L:** Distance of migration, speed of migration, and percentage of migratory phases of Atg5 cKO and WT neuroblasts (n = 188 and 198 cells from 9 Atg5 WT and cKO mice, respectively). *** p < 0.001 with the Student t-test. **M-O:** Example of DAB staining of coronal sections from Atg5 cKO and Atg5 WT mice obtained from brains fixed 7 days after a single tamoxifen injection. **P:** Quantification of GFP+ neuroblast density in the SVZ, RMS, RMSob, and OB of Atg5 WT and cKO mice (n = 5 mice for the Atg5 cKO and WT mice, *p < 0.05). **Q:** Experimental procedure for GFP- or Cre- mNeptune-encoding retroviral labelling of neuroblasts from Atg5 wt/wt and Atg5 fl/fl mice. **R:** Example of time-lapse imaging of neuroblasts expressing GFP or Cre-mNeptune. **S-T:** Distance of migration, speed of migration, and percentage of migratory phases for Atg5 wt/wt and Atg5 fl/fl mice injected with GFP or Cre-mNeptune retroviruses. Atg5 wt/wt-GFP (n = 143 cells from 8 mice), Atg5 wt/wt-Cre-mNeptune (n = 208 cells from 9 mice), Atg5 fl/fl-GFP (n = 106 cells from 7 mice), and Atg5 fl/fl-Cre-mNeptune (n = 121 cells from 5 mice). *** p < 0.001 with a one-way ANOVA followed by an LSD-Fisher post hoc test. **V:** Experimental procedure for the electroporation of plasmids expressing Cas9 and gRNAs. Plasmids were injected into the lateral ventricle of P1-P2 pups followed by the application of electrical pulses. Acute sections were prepared 8-15 days post-electroporation, and the migration of electroporated cells was assessed by time-lapse imaging. **W:** Time-lapse imaging of neuroblasts electroporated with control and LacZ gRNAs in acute brain sections. **X-Z:** Distance of migration, speed of migration, and percentage of migratory phases of cells electroporated with Atg12 gRNAs (n = 19 and 42 cells for LacZ and Atg12 gRNAs, respectively). * p < 0.05 and *** p < 0.001 with Student t-test. Individual values as well as means ± SEM for all time-lapse imaging experiments are shown. See also **Supplementary Figures 1 and 2, and Supplementary Movie 2 and 3**.

We next used the genetic strategy to ablate an essential autophagy gene, Atg5. We thus used the transgenic GlastCreErt2::Atg5fl/fl::GFP mouse model in which a tamoxifen injection causes the inducible conditional knock-out of Atg5 (hereafter, Atg5 cKO) in stem cells and their progeny (**Figure 2G**). We first verified the efficiency of autophagy impairment by performing an EM analysis of GFP+ neuroblasts in the RMS of GlasCreErtT2::Atg5wt/wt::GFP (Atg5 WT) and Atg5 cKO mice. The analysis showed that area of cells occupied by autophagosomes was 75% lower in the neuroblasts of the Atg5 cKO mice without any changes in individual autophagosome size, indicating a decreased in the total numbers of autophagosome (**Supplementary Figure 1**). These results confirmed that impairment of autophagy in Atg5 cKO cells was efficient. Time-lapse imaging of migrating neuroblasts in acute brain sections from Atg5 WT and cKO mice showed a defect in the distance of migration for Atg5-deficient neuroblasts (43.4 ± 1.5 μm in control mice vs. 28.5 ± 1.3 μm; *p < 0.001*, n = 188 and 198 cells from 9 mice per condition, for the Atg5 WT and cKO mice, respectively; **Figure 2G-J, Supplementary Movie 2**). This decrease was caused by a reduction in the percentage of the migratory phases (47.8 ± 0.8% in WT mice vs. 35.1 ± 0.8% in Atg5 cKO mice;*p < 0.001*), with no change in the speed of migration, which was estimated exclusively during the migratory phases (116.2 ± 2.3 μm/h in WT mice vs. 110.4 ± 2.4 μm/h in Atg5 cKO mice; **Figure 2G-L**). To study the impact of autophagy suppression on the total neuroblast population, we performed GFP immunostaining on serial coronal sections from the OB to SVZ of the Atg5 WT and cKO mice 7 days after a single tamoxifen injection (**Figure 2M-P).** An Atg5 deficiency led to the accumulation of neuroblasts in the RMS close to the SVZ (582.7 ± 72.5 cell/mm^2^ in WT mice vs. 846.7 ± 72.7 cell/mm^2^ in cKO mice; *p<0.05*, n = 4 animals per group), with an accompanying decrease in the density of neuroblasts in the rostral RMS (RMS of the OB) and the OB. Taken together, these results show that Atg5-dependant autophagy is required to maintain a normal periodicity of migratory and stationary phases of migration and the correct routing of neuroblasts toward the OB.

In these experiments, the expression of Atg5 was affected at the very beginning of the stem cell lineage, which may have indirectly affected the migration of the progeny derived from these stem cells. We thus stereotactically injected a mixture of retroviruses encoding Cre-mNeptune or GFP into the SVZ of Atg5fl/fl and Atg5 wt/wt mice (**Figure 2Q-U**). The retroviruses infect rapidly dividing cells such as neuroblasts but not slowly dividing stem cells. Atg5fl/fl neuroblasts expressing Cre displayed the same deficiency in migratory parameters as neuroblasts in Atg5 cKO mice (**Figure 2S-U; Supplementary Movie 3**). Importantly, neuroblasts in Atg5 wt/wt mice infected either with Cre-mNeptune or GFP retroviral particles or neuroblasts of Atg5fl/fl mice infected with GFP retroviral particles displayed normal migration patterns (**Figure 2S-U**). These results suggest that the deficiency in cell migration does not arise from the progression and differentiation of the stem cell lineage. It has been recently reported that Atg5 deletion may produce an autophagy-independent effect (Galluzzi and Green, 2019). To verify this possibility, we genetically impaired the expression of Atg12, another essential autophagy-related gene, using CRISPR/Cas9 technology. We electroporated plasmids carrying Cas9-T2A-mCherry and gRNAs against Atg12 in the early postnatal period. We used gRNAs directed against LacZ as a control. We performed time-lapse imaging of mCherry+ cells in the RMS 8-15 days later and observed that Atg12 gRNAs cause the same defects in cell migration, as Atg5 deficiency (**Figure 2V-Z**). Altogether, our results with pharmacological and genetic perturbations of autophagy in the migrating cells indicate the requirement of this self-catabolic pathway for cell migration.

We next asked how autophagy affects cells migration. Previous studies have shown that paxillin, a focal adhesion protein, is a direct target of LC3-II and is recycled by autophagy during migration of non-neuronal cells *in vitro* (Kenific et al., 2016; Sharifi et al., 2016). Since induction of autophagy and cargo recycling is cell- and context-dependent (Klionsky et al., 2016) we thus asked whether paxillin may be also recycled by autophagy in migrating neurobalsts *in vivo*. We electroporated plasmids carrying paxillin-GFP and RFP-LC3 in pups and analyzed their co-localization 7-10 days later in the RMS **(Figure 3A)**. Our analysis revealed a 27.6 ± 1.7 % (n=30 cells from 3 mice) of colocalization of paxillin-GFP in RFP-LC3 vesicles in WT mice **(Figure 3B)**, suggesting for active recycling of paxillin in migrating neuroblasts in the RMS. To assess whether lack of autophagy leads to altered recycling of paxillin, we performed immunolabelling for paxillin in Atg5 WT and cKO mice. We observed that deficiency in autophagy leads to increased immunolabelling of paxillin in the leading process on neuroblasts with no changes at the level of cell soma **(Figure 3 C-D)**. Altogether these data indicate autophagy regulates neuronal migration *in vivo* via recycling of paxillin.

**Figure 3:**
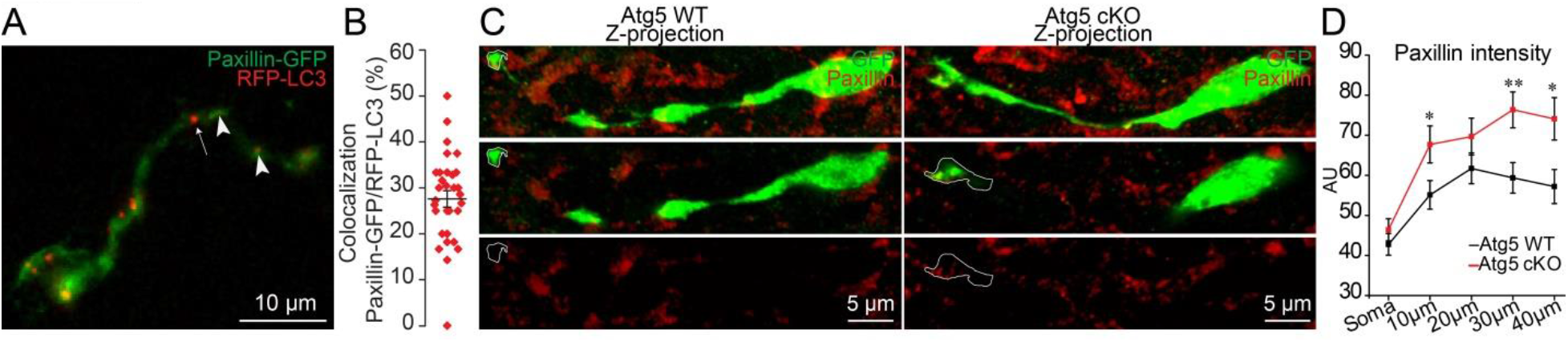
Autophagy affects neuronal migration via recycling of paxillin. A-B: Example and quantification of paxillin-GFP and RFP-LC3 dots in migrating neuroblasts. Arrowheads indicate paxillin-GFP/RFP-LC3 dots, whereas arrow shows RFP-LC3 puncta only. **C**: Example of paxillin immunolabelling in neuroblasts of Atg5 WT and cKO mice. **D**: Quantification of fluorescence intensity of paxillin at the level of cell body and leading process with the bins of 10 μm (n= 38 cells from 3 mice for Atg5 WT and n=31 cells from 3 mice for Atg5 cKO) * p < 0.05 and ** p < 0.005 with the Student t-test.

### Autophagy is induced by bioenergetic requirements during cell migration

Autophagy is associated with energetic stress (Kroemer et al., 2010). The energy status of cells, represented by the ATP/ADP ratio, is closely related to the ability of metastatic cancer cells to migrate *in vitro* (Zanotelli et al., 2018). We used a ratiometric biosensor of ATP/ADP (PercevalHR) (Tantama et al., 2013) to explore energy consumption during cell migration *in vivo* and to investigate the link between energy consumption and autophagy. PercevalHR- and TdTomato-encoding lentiviruses were co-injected into the SVZ of adult mice. Time-lapse imaging of acute sections was performed to assess the changes in the ATP/ADP ratio during the different phases of cell migration in TdTomato-labeled neuroblasts in the RMS (**Figure 4A, Supplementary Movie 4**). Interestingly, the ATP/ADP ratio was dynamically regulated during the different phases of cell migration (**Figure 4B-C**) similar to the dynamic changes observed for autophagosomes (**Figure 1I**). Shortly after the beginning of the migratory phases, the ATP/ADP ratio started to decrease, with a peak ~20-fold decrease. The decrease was accompanied by the entry of the cells into the stationary phase and a progressive recovery of the ATP/ADP ratio (**Figure 4B-C**). These results indicate that the ATP/ADP ratio is dynamically regulated during cell migration and suggest the presence of mechanisms leading to the recovery of ATP levels following the decrease during the migratory phases. The main sensor of energy levels in cells is an AMP-activated kinase (AMPK), which is phosphorylated when the ATP/ADP ratio decreases (Hardie et al., 2012). The phosphorylation of AMPK may induce autophagy (Egan et al., 2011) by phosphorylating Ulk1 and Ulk2 (Klionsky et al., 2016) which may mechanistically link the observed dynamic changes in the autophagosomes and the ATP/ADP ratio during the migratory and stationary phases. To assess the link between energy consumption and autophagy in the migrating cells we used genetic and pharmacological approaches. Since phosphorylation of Ulk1 and Ulk2 by AMPK is crucial for the induction of autophagy, we first electroporated plasmids carrying Cas9-T2A-mCherry and gRNAs against Ulk1 and Ulk2 (and LacZ as a control) in the early postnatal period (**Figure 4D-E**). We observed reduced distance of migration of neuroblasts electroporated with Ulk1 or Ulk 2 gRNAs (**Figures 4E-F**), because of longer stationary phases (**Figure 4H**) without any effect on the speed of migration (**Figure 4G**). To investigate further the interplay between energy consumption and autophagy in the migrating cells, we next used a pharmacological approach to block the AMPK using Compound C (cC) (Kim et al., 2011) in control and Atg5-deficient neuroblasts. We argued that if AMPK induces activation of autophagy, then blocking this kinase by cC in autophagy-deficient neuroblasts should not have any additional effect. Acute sections were prepared from GlastCreErt2::GFP mice following a tamoxifen injection, and the cells were imaged in the presence of cC (**Figure 4I-K**). cC reduced the distance of cell migration because of a decrease in the percentage of the migratory phases but not of the speed of migration (**Figure 4N-P**). We next inhibited AMPK with cC in acute sections from Atg5 cKO mice and observed no additional changes in the distance of migration or in the duration of the migratory phases (**Figure 4L-P**). These results highlight the importance of AMPK in controlling the periodicity of the migratory and stationary phases during cell migration and indicate that changes in the energy status of cells induce autophagy.

**Figure 4:**
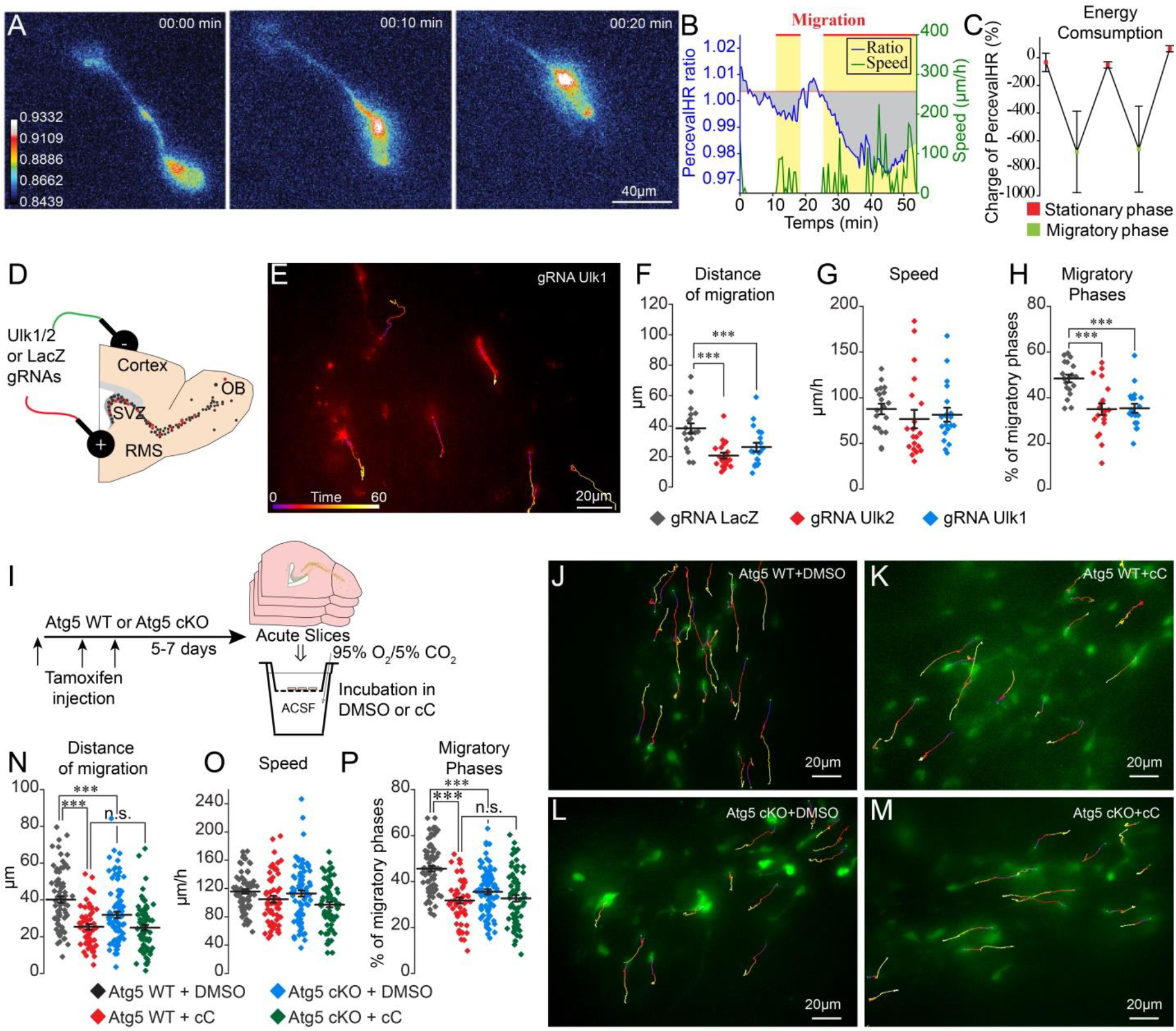
ATP/ADP ratios are dynamically modulated during cell migration and trigger autophagy via AMPK. **A-B:** Example and quantification of changes in the ATP/ADP ratios of individual neuroblasts during different phases of cell migration. Yellow boxes indicate the migratory phases. Gray areas indicate the charge (integrated PercevalHR ratio). **C:** Quantification of changes in the ATP/ADP ratio of neuroblasts during the migratory and stationary phases (n = 19 cells). Data are expressed as means ± SEM. **D:** Experimental procedure for the electroporation of plasmids expressing Cas9 and gRNAs. **E:** Time-lapse imaging of neuroblasts electroporated with Ulk1 gRNAs in acute brain sections. **F-H:** Distance of migration, speed of migration, and percentage of migratory phases of cells electroporated with LacZ, Ulk1 and Ulk2 gRNAs (n = 19 cells for LacZ and Ulk1 gRNAs, and 20 cells for Ulk2 gRNAs, respectively). * p < 0.05 and *** p < 0.001 with a one-way ANOVA followed by an LSD-Fisher post hoc test. **I:** Experiments to study the role of AMPK in autophagy-dependent neuronal migration. Acute sections from Atg5 WT and cKO mice were incubated for 2 h with DMSO or Compound C (cC, 20μM) followed by time-lapse imaging of cell migration in the presence of DMSO or cC. **J-M:** Example of time lapse imaging of cell migration of Atg5 WT and cKO neuroblasts in the presence of DMSO or cC. **N-P:** Distance of migration, speed of migration, and percentage of migratory phases of neuroblasts (n = 79 cells from 5 mice for Atg5 WT+DMSO, n = 56 cells from 5 mice for Atg5 WT+cC, n = 85 cells from 6 mice for Atg5 cKO+DMSO, and n = 72 cells from 7 mice for Atg5 cKO+cC). *** p < 0.001 with a one-way ANOVA followed by an LSD-Fisher post hoc test. Individual values as well as means ± SEM for all time-lapse imaging experiments are shown. See also **Supplementary Movie 4**.

### The pharmacological modulation of cell migration induces changes in autophagy

While our results show that pharmacological and genetic perturbations in autophagy-related genes affect cell migration and that energy levels are linked to autophagy induction, it remains unclear whether autophagy levels are dynamically regulated in response to migration-promoting or inhibiting cues to sustain cell migration. To address this issue, we incubated adult forebrain sections with factors known to affect neuroblasts migration in the RMS (Shinohara et al., 2012; Snapyan et al., 2009; Lee et al., 2006; Bolteus and Bordey, 2004) dissected out the RMS, and immunoblotted these samples for LC3 (**Figure 5**). We used BDNF (10 ng/mL), which is known to promote migration of neuroblasts (Snapyan et al., 2009), GABA (10 μM), which reduces the speed of migration (Snapyan et al., 2009; Bolteus and Bordey, 2004), GM60001 (100 μM), which inhibits matrix metalloproteinases and impact neuroblasts migration in the RMS (Rempe et al., 2018; Lee et al., 2006), Y27632 (50 μM) and blebbistatin (100 μM), which are known to affect neuroblasts migration by inhibiting Rock and myosin II (Shinohara et al., 2012). Intriguingly, all these treatments induced changes in LC3-II, the lipidated autophagosomal form of LC3 (**Figure 5 A,D**), suggesting that changes in autophagy are a common signature for neurons displaying either up- or down-regulated migration. Since increases in LC3-II levels could reflect either an increase in autophagic flux or an accumulation of autophagosomes (Klionsky et al., 2016), we investigated proteins that may be sequestered and recycled by the autophagic process as p62/SQSTM1, a major autophagy substrate (Pankiv et al., 2007), or paxillin (Kenific et al., 2016; Sharifi et al., 2016). We thus performed immunoblotting for p62 and paxillin on RMS samples. Interestingly, while the level of p62 did not change after these pharmacological treatments (**Figure 5B**), the level of paxillin increased when cell migration decreased and decreased when cell migration increased (**Figure 5A-E**). This is in line with our previous analysis showing the presence of paxillin-GFP in RFP+ autophagosomes and accumulation of paxillin in Atg5-deficient neuroblasts (**Figure 3**). To provide further support for dysregulated autophagy in the context of affected cell migration, we immunostained sections containing neuroblasts infected with RFP-GFP-LC3 for Lamp1, a lysosome marker, to determine the autophagosome/autolysosome ratio (**Figure 5F-G**). In line with previous results, drugs that decrease migration led to an increase in the autophagosome/autolysosome ratio, indicating deficient autophagy (**Figure 5G**). These results indicate that several signaling pathways known to affect neuroblasts migration converge on autophagy to sustain the pace and periodicity of cell migration through a constant assembly and disassembly of focal adhesions required for cell motility.

**Figure 5:**
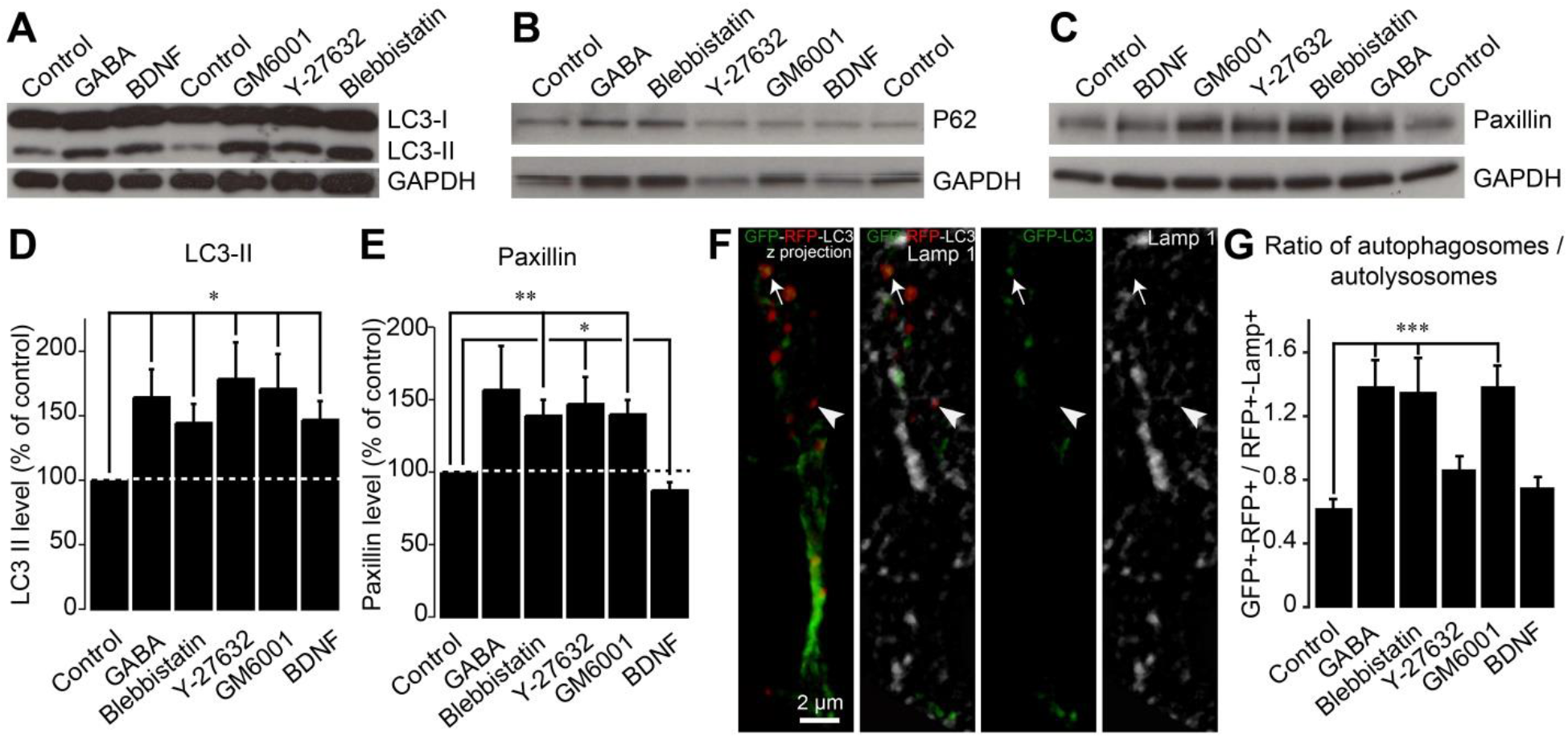
Autophagy is dynamically regulated by migration-promoting and inhibiting cues and is required for the recycling of paxillin. **A-C:** Immunoblotting for a lipidated form of LC3 (LC3-II), p62, and paxillin on RMS samples dissected from acute sections previously incubated with BDNF, GABA, GM60001, Y27632, or blebbistatin for 2 h. GAPDH was used as a housekeeping protein. **D-E:** Quantification of LC3-II and paxillin levels after the pharmacological manipulation of cell migration (n = 5-7 mice for both groups, * p < 0.05 and ** p < 0.005 with a Student t-test). **F**: Example of a cell infected with a retrovirus expressing the LC3-GFP-RFP fusion protein and immunostained for Lamp1 to label autophagosomes (GFP+/RFP+) and autolysosomes (RFP+/Lamp1+). **G:** Percentage of autophagosomes and autolysosomes after a 2-h incubation with BDNF, GABA, GM60001, Y27632, or blebbistatin. Autophagosome/autolysosome ratios was accessed for each cell and the results are expressed as the mean ± SEM. *** p < 0.001 with a one-way ANOVA followed by a post hoc LSD-Fisher test.

## Discussion

We show that cell migration is sustained by a dynamic interplay between ATP/ADP levels and autophagy, which concomitantly regulates the pace and periodicity of the migratory and stationary phases. The pharmacological and genetic impairments of autophagy and AMPK both reduced cell migration by prolonging the stationary phases. Autophagy was also dynamically regulated in response to migration-promoting or inhibiting factors and was required for the recycling of the focal adhesion molecule paxillin.

Tracking intracellular ATP/ADP levels in migrating cells allowed us to demonstrate that the efficiency of cell migration and the periodicity of the migratory and stationary phases are directly linked to the energy status of the cells. The migratory phase led to a drop in ATP/ADP levels that, in turn, was associated with the entry of the cells into the stationary phase. During the stationary periods, ATP/ADP levels recovered due to AMPK activation, following which the cells re-entered the migratory phase. Although it is known that cell migration entails a considerable bioenergetic demand (LeBleu et al., 2014; Zhou et al., 2014; van Horssen et al., 2009), it remains unclear whether and how energy consumption is dynamically regulated during cell migration. Zhang et al. used intracellular measurements of ATP/ADP levels during breast cancer cell invasion to show that energy levels play a role in the coordinated leader-follower cell transition during collective migration (Zhang et al., 2019). ATP/ADP levels are also proportional to the propensity of cancer cells to migrate *in vitro* (Zanotelli et al., 2018). Our results further strengthen the link between cell migration and ATP/ADP levels and show that dynamic changes in ATP/ADP levels are required for sustaining cell migration by regulating the duration and periodicity of the migratory and stationary phases.

We also show that a decrease in ATP/ADP levels leads to the activation of AMPK, which in turn triggers autophagy. One of the downstream targets of AMPK is Ulk1/2 (Egan et al., 2011) and Ulk1/FIP2000 complex is a key activator of autophagy (Jung et al., 2009; Hara et al., 2008). This may provide a mechanistic link between energy consumption and autophagy induction during cell migration. Indeed, the genetic impairment of Ulk1, Ulk2, and other autophagy genes such as Atg5 and Atg12 in neuroblasts resulted in impaired cell migration, while the inhibition of AMPK in Atg5-deficient cells failed to induce any additional changes in cell migration. The present study thus adds to the growing body of evidence indicating that autophagy plays a major role in cell migration. Previous studies were, however, performed *in vitro* using non-neuronal cells and resulted in conflicting results (Kenific et al., 2016; Sharifi et al., 2016; Li et al., 2015; Tuloup-Minguez et al., 2013). shRNA against Atg7 or a knock-out of Atg5 increases the migration of HeLa cells and murine embryonic fibroblasts (Tuloup-Minguez et al., 2013) and endothelial progenitor cells (Li et al., 2015). In contrast, it has been shown that autophagy promotes cell migration by recycling focal adhesions while the genetic impairment of autophagy leads to decreased cell migration (Kenific et al., 2016; Sharifi et al., 2016). Since autophagy depends on the cellular context (Klionsky et al., 2016), it is conceivable that differences in these studies can be explained by the types of cells studied and the cellular context in which they were studied. Our *ex vivo* and *in vivo* results show that autophagy promotes migration of developing neurons. Furthermore, we show that autophagy is very dynamic during cell migration, is dynamically regulated in response to migration-promoting and inhibiting factors, and is involved in the recycling of focal adhesions. These results are in agreement with those of previous *in vitro* studies showing that LC3-II directly targets the focal adhesion protein paxillin (Kenific et al., 2016; Sharifi et al., 2016). Changes in autophagy levels in response to migration-promoting or inhibiting factors are required to cope with the capacity of cell to migrate by the recycling of focal adhesions. Indeed, our results suggest that decreases in cell migration are linked to a lower recycling rate of paxillin and a higher autophagosome/autolysosome ratio, while the promotion of cell migration tends to increase paxillin turnover. During cell migration, autophagy not only recycles focal adhesion proteins, but also inhibits focal adhesion kinase through the Atg13/Ulk1/FIP2000 complex, which promotes cell immobility (Caino et al., 2013). Autophagy activator protein VPS15, which is part of VPS34-PI3-kinase complex I, also mediates the stability of the cytoskeleton proteins actin and tubulin by decreasing Pak1 activity (Gstrein et al., 2018). On the other hand, it has also been shown that autophagy is involved in the secretion of matrix metalloproteases (MMPs) and pro-migratory cytokine interleukin-6 (IL6), which promotes tumor cell migration (Lock et al., 2014). These results suggest that autophagy may play a dual role in controlling cell migration. First, it mediates cytoskeleton stability, which maintains the stationary phase, which is required for cellular homeostasis and the regeneration of ATP/ADP levels. Second, it mediates the recycling of focal adhesions and the release of pro-migratory factors, which are required for the entry of cells into the migratory phase. Altogether, our results reveal that the dynamic interplay between energy levels and autophagy in migrating cells is required to sustain neuronal migration and the periodicity of the migratory and stationary phases.

## Materials and Methods

### Animals

Experiments were performed using two- to four-month-old C57BL/6 mice (Charles River, strain code: 027), glial fibrillary acidic protein (GFAP)-GFP mice (The Jackson Laboratory, strain code: 003257FVB/N-Tg(GFAPGFP)14Mes/J), and GlasCreERT2::Atg5fl/fl::CAG-CAT-GFP (Atg5 cKO, strain code: B6.129S-Atg5<tm1Myok>, RRID:IMSR_RBRC02975) and GlasCre-ERT2::Atg5wt/wt::CAG-CAT-GFP (Atg5 WT) mice as well as two-week-old CD-1 pups (Jackson laboratories, strain code: 022), which were electroporated on postnatal day 1-2. To obtain the Atg5 WT and cKO mice, we first crossed GlastCreErt2 mice (Mori et al., 2006) with CAG-CAT-GFP mice (Waclaw et al., 2006) to obtain GlastCreErt2::GFP mice. This mouse strain was then crossed with Atg5fl/fl mice (B6.129S-Atg5<tm1Myok>, RBRC02975, Riken) to obtain Atg5 WT and cKO mice. All the experiments were approved by the Université Laval animal protection committee. The mice were housed one to five per cage. They were kept on a 12-h light/dark cycle at a constant temperature (22°C) with food and water *ad libitum*.

### Stereotactic injections

For retro- or lentivirus injections, C57BL/6 mice were anesthetized with isoflurane (2-2.5% isoflurane, 1 L/min of oxygen) and were kept on a heating pad during the entire surgical procedure. Lentiviruses or retroviruses were stereotactically injected in the dorsal and ventral subventricular zones (SVZd and SVZv) at the following coordinates for C57BL/6 (with respect to the bregma): anterior-posterior (AP) 0.70 mm, medio-lateral (ML) 1.20 mm, dorso-ventral (DV) 1.90 mm, and AP 0.90 mm, ML 1 mm, and DV 2.75 mm. For Atg5 fl/fl and Atg5 wt/wt following coordinates were used: AP 0.90 mm and AP 1.10 mm was used respectively for SVZd and SVZv. The following viruses were used: CMV-PercevalHR (1×10^10^ TU/mL, produced at the University of North Carolina Vector Core Facility based on plasmid # 49083, Addgene, kindly provided by Dr. Yellen, Harvard Medical School), CMV-Td-Tomato-encoding lentivirus (1.5×10^10^ TU/mL, produced at the University of North Carolina Vector Core Facility based on plasmid # 30530, Addgene, kindly provided by Dr. Ryffel, University of Duisburg-Essen), RFP-GFP-LC3-encoding retrovirus (9.3×10 TU/mL, produced at the Molecular Tools Platform at the CERVO Brain Research Center based on plasmid #21074, Addgene, kindly provided by Dr. Yoshimori, Osaka University), GFP-encoding retrovirus (2.9×10 TU/mL, Molecular Tools Platform at the CERVO Brain Research Center), and Cre-mNeptune-encoding retrovirus (4×10^7^ TU/mL, Molecular Tools Platform at the CERVO Brain Research Center).

For the electroporation of plasmids, P1-P2 CD-1 pups were anesthetized using isoflurane (2-2.5% isoflurane, 1 L/min of oxygen). The plasmids (1.3 μL, 3-6 μg/μL total) were injected in the lateral ventricle using the following coordinates (with respect to the lambda): AP 1.8 mm, ML 0.8 mm, and DV 1.6 mm. Immediately after the injection, an electric field (five 50-ms pulses at 100 mV at 950-ms intervals) was applied using an electrode positioned on the surface of the bones. The pups were used 8 to 15 days post-injection. We used gRNAs for Ulk1, Ulk2, and Atg12. We used gRNAs against LacZ as a control. We used two different gRNAs for each gene and electroporated these plasmids together. To assess the autophagy-dependent recycling of paxillin, we co-electroporated pmRFP-LC3 (Addgene, #21075, kindly provided by Dr. Yoshimori, Osaka University) with pRK paxillin-GFP (Addgene, #50529, kindly provided by Dr. Yamada, National Institute of Dental and Craniofacil Research) or paxillin-GFP (kindly provided by Dr. Giannone, Université Bordeaux Segalen) at P1-P2 and analyze migrating neuroblasts in the RMS 8-10 days later.

### CRISPR target site selection and assembly

The gRNAs were designed and selected using ChopChop online software (Labun et al., 2019). We used two different gRNAs to target each gene to increase the efficiency of the CRISPR editing. The gRNAs were cloned in the BbsI site of the PU6-BbsI-CBh-Cas9-T2AmCherry plasmid (Addgene, #64324, kindly provided by Dr. Kuehn, Berlin Institute of Health). The following gRNA sequences were used:

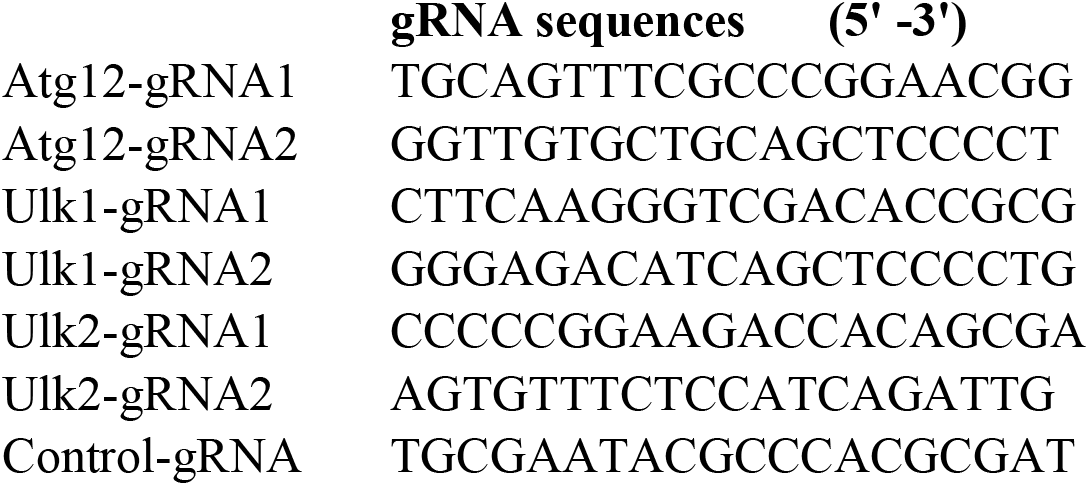

The efficiency of the gRNAs was verified by high-resolution melting curve PCR (Thomas et al., 2014) on primary cultures of adult NPCs transfected with above-mentioned plasmids. Briefly, the thin layer of SVZ bordering the lateral ventricle, excluding the striatal parenchyma and corpus callosum, was dissected from the SVZ of 8- to 12-week-old C57BL/6 mice. The tissue was minced and digested in a 0.05% trypsin-EDTA solution, following which an equal volume of soybean trypsin inhibitor was added. After trituration, the resulting single cell suspension was cultured in NeuroCult Basal Medium (StemCell Technologies, #05701) with NeuroCult Proliferation Supplement (StemCell Technologies, #05701), EGF and bFGF (10 ng/mL each, Sigma-Aldrich, #E4127 and #SRP4038-50UG), and heparin (2 μg/mL, StemCell Technologies, #07980).

Genomic DNA was isolated using DNeasy Blood & Tissue kits (QIAGEN, #69504) according to the manufacturer’s protocol. Primers were designed using Primer-Blast software (https://www.ncbi.nlm.nih.gov/tools/primer-blast/).

The PCR reactions were performed with 5 μL of LightCycler^®^ 480 High Resolution Melting Master (Roche, #04909631001), 0.5 μL of each primer (10 μM), 1.2 μL of MgCl_2_ (25 mM, Sigma-Aldrich, #M8266; CAS 7786-30-3), 2 μL of genomic DNA, and completed by water to 10 μL. The PCR was performed in a LightCycler 480 (Roche) using 96 well plates (Bio-Rad). The amplification started with an initial denaturation step at 95°C for 5 min, followed by 48 cycles at 95°C for 10 s, 60°C for 30 s, and 72°C for 25 s. Melting curves were generated over a 65-95 °C range in 0.2°C increments and were analyzed using LightCycler 480 SW1.5.1 software. The following primers were used:

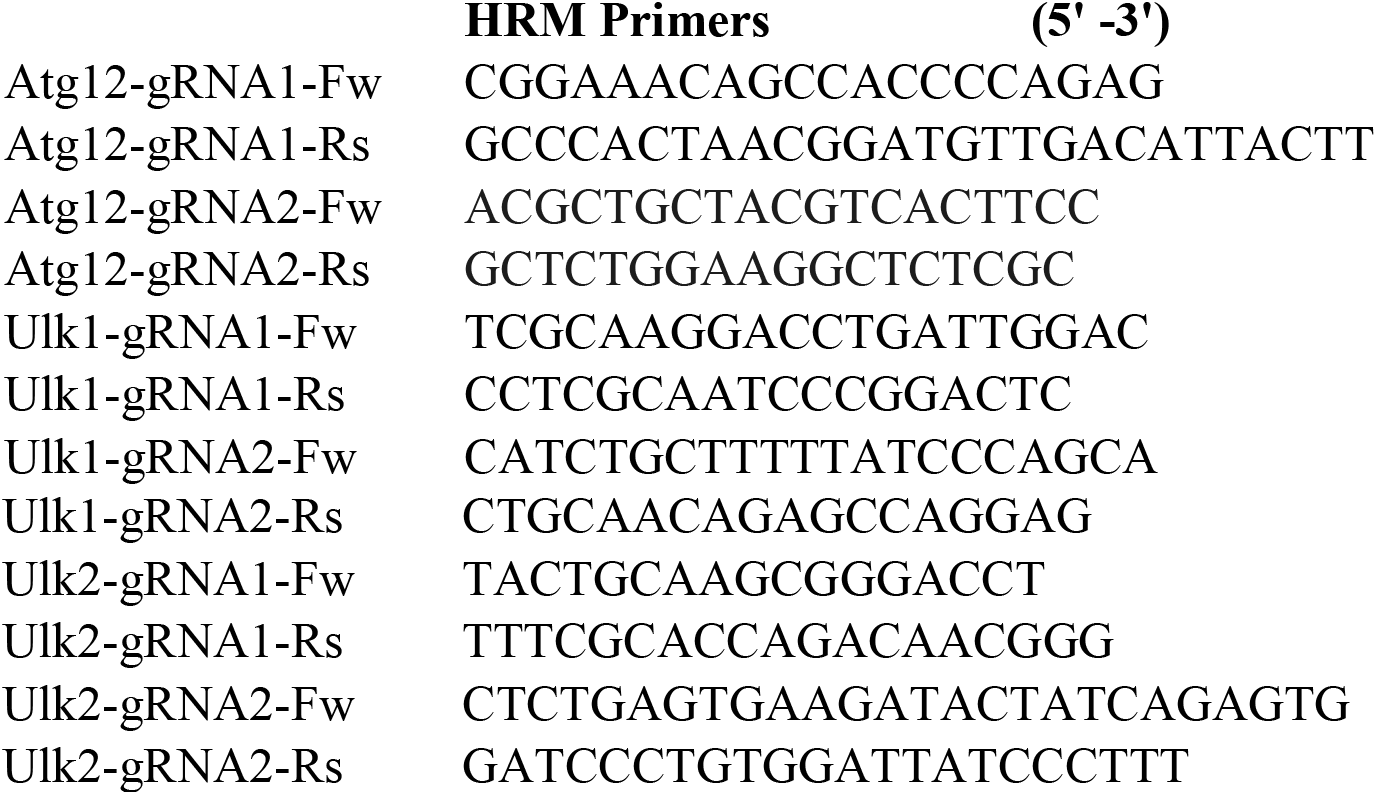

### Tamoxifen injection

For time-lapse imaging of neuroblasts in acute brain sections from Atg5 WT and cKO mice, the tamoxifen was injected intraperitonially (180 mg/kg, Sigma Aldrich, #T5648, CAS: 10540-29-1) once a day, for 3 days. The tamoxifen was diluted in sunflower seed oil (Sigma Aldrich, #S5007, CAS: 8001-21-6) and anhydrous ethanol (10% final, Commercial Alcohols, #1019C). The mice were used 7 to 15 days after the last tamoxifen injection.

The mice received a single intraperitoneal tamoxifen injection (180 mg/kg, Sigma Aldrich) to assess neuroblast distribution *in vivo* along the SVZ-OB pathway. They were sacrificed 7 days later.

### Immunostaining

Animals were deeply anesthetized with sodium pentobarbital (12 mg/mL; 0.1 mL per 10 g of body weight) and were perfused intracardially with 0.9% NaCl followed by 4% paraformaldehyde (PFA) (Sigma-Aldrich, #P6148; CAS: 30525-89-4). Brains were collected and were kept overnight in 4% PFA. Sagittal sections (30 or 40 μm) were cut using a vibratome (Leica). For immunolabelling experiments with acute brain sections, 250-μm-thick sections were fixed in 4% PFA overnight and were pre-permeabilized with methanol and acetone (30 min each at −30°C) prior to immunostaining. The brain sections were incubated with the following primary antibodies: anti-LC3B (1:200, Novus, #NB100-2220, RRID:AB_10003146), anti-LC3A (1:100, Abgent, #AP1805a, RRID:AB_2137587), anti-Atg5 (1:400, Novus, #NB110-53818, RRID:AB_828587), anti-Lamp1 (1:500, EMD Millipore, #AB2971, RRID:AB_10807184), and anti-GFP (1:1000, Avés, #GFP-1020; RRID:AB_1000024). For anti-LC3A, LC3B and Atg5, sections were incubated in 10 mM citrate buffer (pH 6.0) for 20min at 80°C. The primary antibodies were diluted in either 0.2% or 0.5% Triton X-100 and 4% milk. Images were acquired using an inverted Zeiss microscope (LSM 700, AxioObserver) with a 20X air immersion objective (NA: 0.9) or a 63X oil immersion objective (NA: 1.4).

For paxillin immunolabelling experiments, the brains were kept in 30% sucrose solution for overnight and 15μm thick cryostat sections were prepared. The sections were incubated with anti-paxillin (1:100, BD biosciences, #610051, RRID:AB_397463) and anti-GFP (1:1000, Avés) primary antibodies diluted in 0.2% Triton X-100 and 4% BSA. Images were acquired using an inverted Zeiss microscope (LSM 700, AxioObserver) with a 63X oil immersion objective (NA: 1.4) and fluorescence intensity was assessed using ImageJ after application of background subtraction (50 pixels rolling ball radius) and median filter (2 pixels radius) plugins. Mean fluorescence intensity was calculated for cell soma and leading process of GFP+ neuroblasts in the RMS of Atg5 WT and cKO mice with the bins of 10 μm.

For diaminobenzidine (DAB) staining, 40-μm coronal sections separated by 240 μm were taken from the anterior tip of the OB to the end of the SVZ (approximately at the bregma) of Atg5 WT and cKO mice. They were incubated in 3% H_2_O_2_ prior and then with an anti-GFP primary antibody (1:1000, Thermofisher, #A-11122; RRID:AB_221569) The sections were subsequently incubated for 3 h at room temperature with a biotinylated anti-rabbit IgG antibody (1:1000, EMD Millipore, #AP132B, RRID:AB_11212148) followed by 1 h with avidin-biotin complex (ABC Vector Laboratories, #PK4000) and then with 0.05% 3,3-diaminobenzidine tetrahydrochloride (DAB) and 0.027% H_2_O_2_ in 1 M Tris-HCl buffer (pH 7.6). They were dehydrated in graded ethanol baths and were mounted and coverslipped in DePeX (VWR).

### Electron microscopy

Two weeks after the i.p. injection of tamoxifen, 7 mice (4 Atg5 WT and 3 cKO) were deeply anesthetized with a mixture of ketamine (100 mg/kg, i.p.) and xylazine (10 mg/kg, i.p.). They were transcardially perfused with 40 mL of ice-cold sodium phosphate-buffered saline (0,1 M PBS; pH 7.4), followed by 100 mL of cold 2% acroleine and 100 mL of cold 4% PFA to which 0.1% glutaraldehyde was added. The brains were excised and were post-fixed for 4 h in 4% PFA at 4°C. They were then cut with a vibratome (model VT1200 S; Leica, Germany) into 50-μm-thick sagittal sections collected in PBS.

After a 30-min incubation in a 0.1 M sodium borohydride/PBS solution at room temperature, the sagittal sections containing the RMS were immunostained for GFP. Briefly, free-floating sections were sequentially incubated at room temperature (RT) in blocking solution containing 2% normal goat serum and 0.5% gelatin (1 h), the same blocking solution containing a rabbit anti-GFP antibody (1:1000, Abcam, #ab290, RRID:AB_303395) (24 h, RT), and then the same blocking solution containing a 1:20 dilution of Nanogold-Fab goat anti-rabbit antibody (Nanoprobe, #2004, RRID:AB_2631182) (24 h, 4°C). After thoroughly rinsing the sections in 3% sodium acetate buffer (pH 7.0), the gold immunostaining was amplified using an HQ Silver enhancement kit (Nanoprobe, # 2012) according to the manufacturer’s protocol. The sections were then osmicated, dehydrated in ethanol and propylene oxide, and flat-embedded in Durcupan (Fluka). Quadrangular pieces containing RMS were cut from the flat-embedded GFP-immunostained sections. After being glued to the tip of a resin block, 50-nm sections were cut using an ultramicrotome (model EM UC7, Leica). The ultrathin sections were collected on bare 150-mesh copper grids, stained with lead citrate, and examined with a transmission electron microscope (Tecnai 12; Philips Electronic, 100 kV) equipped with an integrated digital camera (XR-41, Advanced Microscopy Techniques Corp.). Labeled cell bodies were then randomly selected, and autophagosomal vesicles were identified and measured based on careful selection criteria (Eskelinen, 2008).

### Time-lapse imaging

#### Measurement of the ATP/ADP ratio

Seven days after co-injecting lentiviral particles expressing PercevalHR or TdTomato in the SVZ, the mice were sacrificed and acute sections were prepared as described previously (Bakhshetyan and Saghatelyan, 2015). Briefly, the mice were anesthetized with ketamine (100 mg/kg) and xylazine (10 mg/kg) and were perfused transcardially with modified oxygenated artificial cerebrospinal fluid (ACSF) containing (in mM): 210.3 sucrose, 3 KCl, 2 CaCl_2_.2H_2_O, 1.3 MgCl_2_.6H_2_O, 26 NaHCO_3_, 1.25 NaH_2_PO_4_.H_2_O, and 20 glucose. The brains were then quickly removed, and 250-μm-thick sections were cut using a vibratome (HM 650V; Thermo Scientific). The sections were kept at 37°C in ACSF containing (in mM): 125 NaCl, 3 KCl, 2 CaCl_2_.2H_2_O, 1.3 MgCl_2_.6H_2_O, 26 NaHCO_3_, 1.25 NaH_2_PO_4_.H_2_O, and 20 glucose under oxygenation for no more than 6-8 h. PercevalHR imaging was performed using a BX61WI (Olympus) upright microscope with a 60X water immersion objective (NA = 0.9), a CCD camera (CoolSnap HQ), and a DG-4 illumination system (Sutter Instrument) equipped with a Xenon lamp for rapid wavelength switching. The field of view was chosen to have a sparse virally-labeled cells, and images at 430 nm and 488 nm were acquired. The imaging was performed every 30 s for 2 h with multiple z stacks (4-8 stacks, depending on the cell orientation, at 3-μm intervals). The images were analyzed using a custom script written in Matlab. Briefly, a maximum intensity projection of the z stack was created for each wavelength. The migration of the cells was assessed based on the morphological marker TdTomato. The means of the fluorescence intensities at 430 nm and 488 nm were extracted, and the ratio of fluorescence for these two wavelengths was calculated. The speed of migration, as well as the migratory and stationary periods, were determined as described below. To estimate changes in the ATP/ADP ratio, we measured the charge, which was defined as the change in the ratio of fluorescence for the two wavelengths. The mean of the PercevalHR charge for the stationary phases during the entire time-lapse movie of an individual cell was calculated. The changes in the ATP/ADP ratio during the migratory phases were then normalized to those of the stationary phases. Since the magnitude of the ATP/ADP ratio depends on the duration of the migratory and stationary phases, we also normalized it by dividing it by the duration of the phases.

#### Cell migration analysis

Acute sections from Atg5 WT and cKO mice, as well as from C57BL/6 mice injected with retroviruses and CD-1 pups electroporated with plasmids, were obtained as described previously, and time-lapse imaging was performed using the same imaging system using a mercury arc lamp as the illumination source (Olympus). For the pharmacological manipulations of cell migration, the sections were incubated at 37°C in oxygenated ACSF. The following drugs were used: Y-27632 (50 μM, Cayman Chemical, #10005583-5, CAS: 129830-38-2), blebbistatin (100 μM, Toronto Research Chemicals, #B592490-10, CAS: 674289-55-5), GM6001 (100 μM, Abmole, #M2147-10MG, CAS: 142880-36-2), BDNF (10 nM, Peprotech, #450-02), GABA (10 μM, Tocris, #A5835-25G, CAS: 56-12-2), bafilomycin (4 μM, Cayman Chemical, #11038, CAS: 88899-55-2), and Compound C (20 μM, EMD Millipore, #A5835-25G, CAS: 56-12-2). cC was added to the ASCF during the imaging period for the time-lapse imaging of cell migration following AMPK inhibition. Images were acquired with a 40X objective every 15-30 s for 1 h with multiple z stacks (11 stacks with 3-μm intervals). For assessment of autophagosome dynamic in migrating cells, 5-10 mins stream acquisition with 1.4 to 4.95 frames per second was performed. The tracking analysis was performed using Imaris8 software (Biplane) in order to extract the overall cell displacement and vector of cell migration. We did not take into consideration cells that remains stationary during the entire imaging periods, which likely underestimate the autophagy-dependent effects on cell migration. The distance of migration of cells corresponds to the size of the vector displacement between the coordinates of the first and last position of the cell during the one hour of imaging. Then, instantaneous speed (speed between two consecutive times points) profile of each individual cell was plotted using Origin software, and the migratory phases were determined manually based on the speed in order to assess the duration and periodicity of the migratory and stationary phases. The percentage of migration was calculated as the sum of duration of all migratory events during one hour of imaging. The speed of migration was determined only during the migratory phases).

For autophagosome density analysis during migratory and stationary period, RFP-GFP-LC3 expressing cell were imaged as described above to determine migratory and stationary phases. Autophagosome density (RFP+ puncta) was assessed on separates time points corresponding to the beginning, middle and end of each migratory and stationary phase. The mean density of autophagosomes per migratory and stationary phases was then calculated.

### Western blotting

The acute brain sections of adult mouse forebrain were incubated with the pharmacological compounds. The RMS was manually dissected under an inverted microscope (Olympus FV1000, 10X objective, NA=0.4). The tissue was snap frozen in liquid nitrogen and was then incubated in lysis buffer (50 mM HCl, 1 mM EDTA, 1 mM EGTA, 1 mM sodium orthovanadate, 50 mM sodium fluoride, 5 mM sodium pyrophosphate, 10 mM sodium β-glycerophosphate, 0.1% 2-mercaptoethanol, 1% Triton X-100, pH 7.5) supplemented with a protease inhibitor cocktail III (Calbiochem, #539134). The concentration of total protein was measured using the Bradford assay (BioRad). Proteins were separated on 16% NuPage gels (Invitrogen) in SDS running buffer and were transferred to nitrocellulose membranes (Life Technologies). The following primary antibodies were used: anti-LC3B (1:1000, Novus), anti-p62 (1:500, Proteintech, #18420-1-AP, RRID:AB_10694431), anti-paxillin (1:1000, BD biosciences, #610051, RRID:AB_397463), and anti-GAPDH (1:5000, ThermoFisher, #MA5-15738, RRID:AB_10977387).

### Statistical analysis

Data are expressed as means ± SEM. Statistical significance was determined using an unpaired two-sided Student’s *t*-test or a one-way ANOVA followed by an LSD-Fisher post hoc test, depending on the experiment, as indicated. Equality of variance for the unpaired t-test was verified using the F-test. The levels of significance were as follows: * *p* < 0.05, ** *p* < 0.01, *** *p* < 0.001.

## Acknowledgments

This work was supported by Canadian Institute of Health Research (CIHR) grant PJT-159733 to AS. The authors declare no conflict of interests.

## Author contributions

C.B. and A.S. designed the study. C.B., A.P., and M.S. performed the experiments and analyzed the data. D.G. and M.P. performed the EM analysis. S.L. and P.D.K provided the MATLAB script for the ATP/ADP and autophagy flux analyses. A.S. supervised the project. C.B and A.S wrote the manuscript, taking into consideration the comments of the other authors.

## Legends for Supplementary Figures and Movies

**Movie 1: Time-lapse imaging of LC3-GFP-RFP in migrating neuroblasts**

Time-lapse imaging of RFP+ puncta in neuroblasts. The time is indicated in the upper left corner.

**Movie 2: Neuronal migration of Atg5 WT and cKO neuroblasts in acute sections of the adult RMS**

Time-lapse imaging of GFP+ neuroblasts in sections from Atg5 WT (left) and cKO (right) mice. The time is indicated in the upper left corner.

**Movie 3: Neuronal migration of Cre-Neptune and GFP infected cells in Atg5 fl/fl mice**

Example of time lapse imaging of neuroblasts on acute slices from Atg5 fl/fl mice infected with retroviruses expressing GFP and Cre-mNeptune. The images were acquired every 15 s for 1 h. The time is indicated in the upper left corner.

**Movie 4: The ATP/ADP ratio is dynamically modulated during cell migration**

Time-lapse imaging of neuroblasts infected with PercevalHR-encoding and Td-Tomato-encoding lentiviruses. PercevalHR was used for ratiometric measurements of changes in the ATP/ADP ratio. The ATP/ADP ratio is shown. Note that the ATP/ADP ratio dynamically changes during the different cell migration phases. The time is indicated in the upper left corner.

---

**Supplementary Figure 1.**
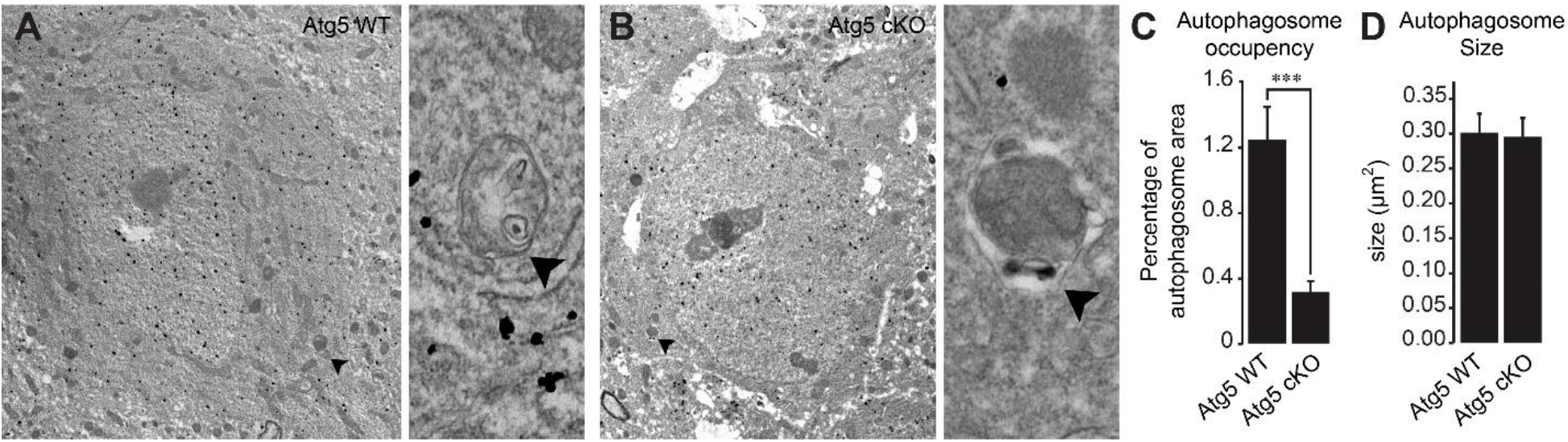
Validation of autophagy impairment in Atg5 cKO mice by electron microscopy. (A-B) Example of electron microscopic images of neuroblasts in the RMS of Atg5 WT and Atg5 cKO mice. The neuroblasts are labeled with anti-GFP immunogold particles. The autophagosome is indicated with an arrowhead. (C) Quantification of the percentage of autophagosome area in Atg5 WT and Atg5 cKO mice. *** p < 0.001 with the Student t-test.

**Supplementary Figure 2.**
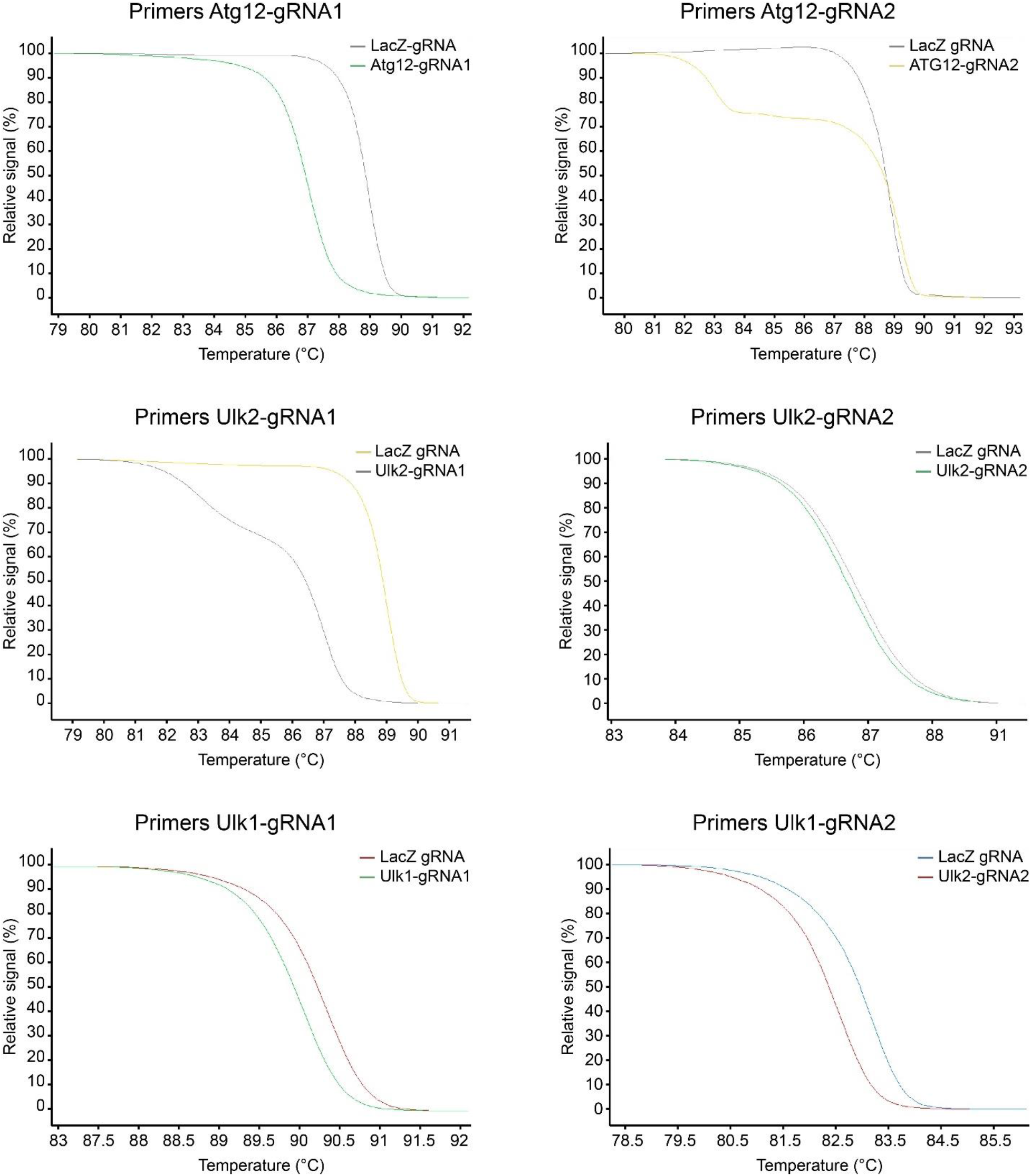
Validation of gRNA efficiency by high-resolution melting (HRM) PCR. SVZ cells were isolated and were cultured *in vitro*. The cells were transfected with plasmids carrying Cas9 and various gRNAs. The PCR reaction was performed on genomic DNA, and HRM curves were generated over a 65-95 °C range in 0.2°C increments.

